# Species-specific, multifaceted venom resistance in *Monodelphis domestica* reveals novel physiological behavior of von Willebrand Factor under flow

**DOI:** 10.1101/2025.01.21.634112

**Authors:** Matthew Holding, Dante Disharoon, Laura Haynes, Bipin Chakravarthy Paruchuri, M. Hao Hao Pontius, Krista Golden, Jordan A. Shavit, Karl Desch, David Ginsburg, Anirban Sen Gupta, Yolanda Cruz, Danielle H. Drabeck

**Affiliations:** Department of Ecology, Evolution, and Behavior, University of Minnesota, Twin Cities; Life Sciences Institute, University of Michigan, Ann Arbor, MI; Department of Ecology and Evolutionary Biology, University of Michigan, Ann Arbor, MI, USA; Department of Biomedical Engineering, Case Western Reserve University, Cleveland, OH 44106, USA; Oberlin College Department of Biology, Oberlin, OH 44074, USA; Department of Pediatrics, University of Michigan, Ann Arbor, MI, USA; Departments of Human Genetics, University of Michigan, Ann Arbor, MI, USA; Departments of Internal Medicine, University of Michigan, Ann Arbor, MI, US

## Abstract

Interactions between predators and prey are often characterized by strong selection pressures that shape extreme physiological adaptations. Venom resistance in large-bodied South American opossums (Clade Didelphini) is a striking example, as these marsupials prey on venomous snakes and exhibit remarkable resistance to their venom. While resistance is well documented in Didelphini, relatively little is known about venom resistance in the smaller, more diverse members of Didelphidae, which inhabit the same regions and encounter the same predators. Here, we investigate venom resistance in the small-bodied opossum, *Monodelphis domestica*, through multi-level physiological assays, examining responses to purified venom components and whole venom from sympatric and allopatric vipers. Our results show *M. domestica* resists venom-induced disruptions to blood coagulation, retains platelet function in the presence of platelet-disrupting venoms, and inhibits snake venom metalloproteinases. Unexpectedly, we find that *M. domestica* von Willebrand Factor (VWF) requires increased shear force to elongate, a previously unknown aspect of opossum blood physiology that may contribute to venom resistance and may have relevance to human coagulopathies. These findings expand the extent of venom resistance beyond large-bodied Didelphini, suggesting it is a widespread trait in South American marsupials and providing new insights into venom-mammal coevolution.

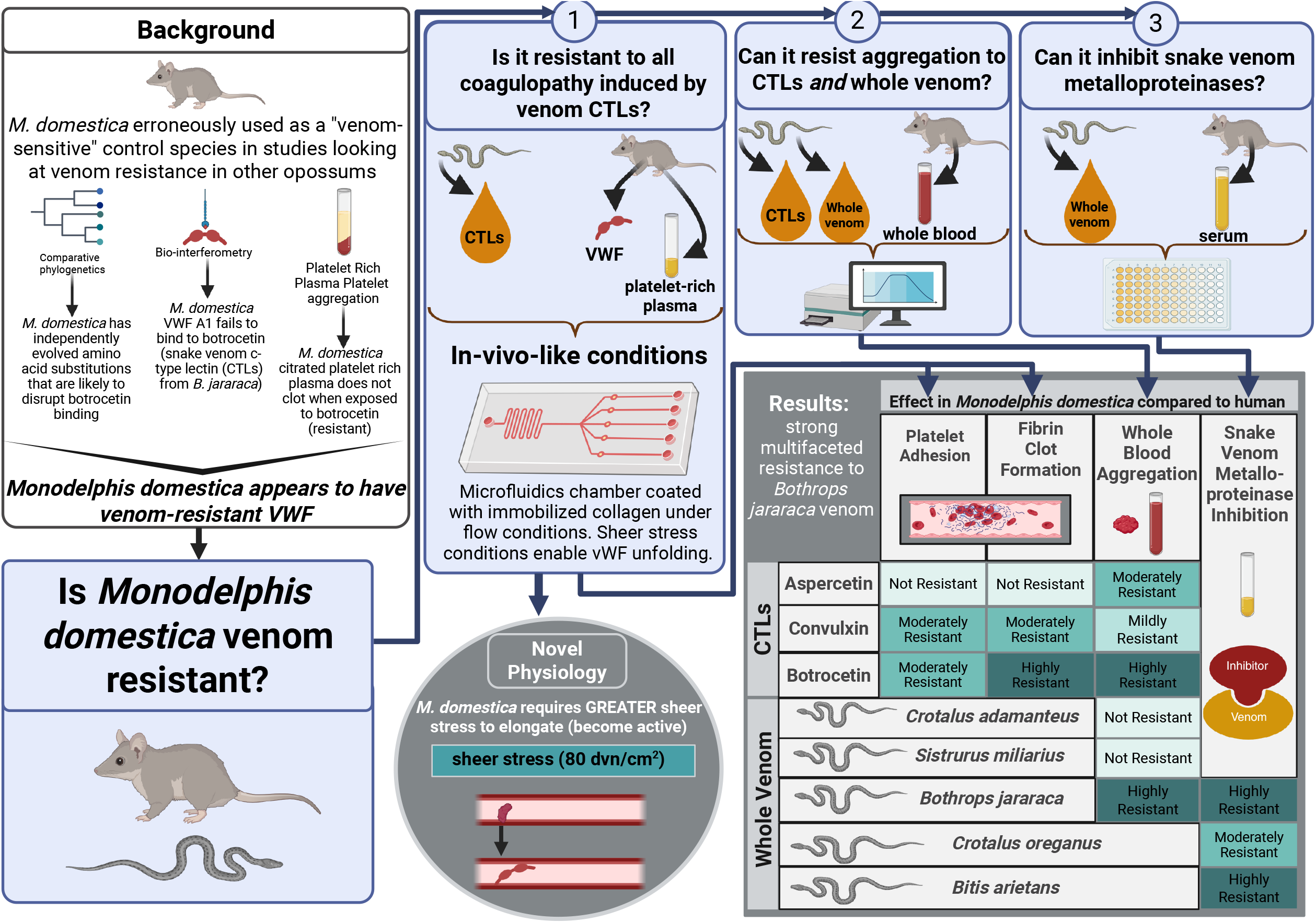

## Introduction

Coevolution between enemies can drive dramatic molecular innovations, producing traits that depart radically from the physiological norms defined in model species like mice and humans (Vermeij 1994; Rowe et al. 2013). When these evolved traits overlap with human-relevant pathways—such as hemostasis—they can yield not only evolutionary insight but unanticipated translational value (King 2011). Venomous species and their prey offer some of the clearest examples of coevolutionary escalation, where reciprocal adaptations emerge even at deeply conserved molecular targets (Holding et al. 2016b; McCabe and Mackessy 2017).

Venomous snakes and their mammalian predators provide one of the clearest natural laboratories for studying coevolution. In South America, members of the opossum clade Didelphini are well-known to hunt and consume pit vipers, including those from the genus *Bothrops*—the continent’s most abundant and widespread group of venomous snakes (Werner and Vick 1977; Voss and Jansa 2012; Jansa and Voss 2011). Large-bodied opossums in this group exhibit remarkable venom resistance: they survive bites that would otherwise be fatal, show minimal coagulopathic symptoms, and have evolved molecular defenses targeting venom components (Drabeck et al. 2020, 2022).

A key component of this resistance involves von Willebrand Factor (VWF), a critical blood-clotting protein that normally mediates platelet adhesion at sites of vascular injury—but is directly targeted by *Bothrops* venom to induce coagulopathy. While many venom CTLs with varying function are known, only botrocetin, bitiscetin, and aspercetin have been identified to specifically target VWF as a coagulopathic intermediary, binding to VWF’s A1 domain and triggering platelet aggregation (Matsushita et al. 2000; Rucavado et al. 2001; Fukuda et al. 2002). The resulting VWF–CTL–platelet complexes are rapidly cleared from circulation, depleting both VWF and platelets and leading to fatal hemorrhage (Bury et al. 2019; Thomazini et al. 2021). This mechanism is further exacerbated by snake venom metalloproteinases (SVMPs), phospholipases, and many other venom components that break down vascular integrity, compounding coagulopathic damage.

In venom-resistant large-bodied opossums, VWF binds poorly—or not at all—to venom-derived CTLs like botrocetin, suggesting that specific mutations on opossum VWF A1 at the botrocetin binding site disrupt this toxin–target interaction and contribute directly to resistance (Drabeck et al. 2022). Surprisingly, VWF from several small-bodied opossum species also fails to bind botrocetin in protein-binding assays, and limited functional data from one small-bodied species (*Monodelphis domestica*) shows reduced platelet aggregation in static assays—despite no prior ecological or physiological evidence of venom resistance. These findings suggest that small-bodied species share venom CTL resistance, but it remains unclear whether this reflects integrated, organism-level resistance to snake venom. While adaptive mutations under positive selection may point to coevolved traits, resistance at a single molecular interface does not establish a complex phenotype. Because venom resistance involves coordinated defenses against multiple toxin classes (including CTLs and SVMPs) further physiological evidence is needed to determine whether small-bodied opossums possess true systemic resistance.

Additionally, while VWF–botrocetin binding assays confirm the molecular basis of resistance in large-bodied opossums (consistent with preserved hemostasis and survival) they are less conclusive in small-bodied species. The absence of detectable binding in vitro does not guarantee functional resistance in vivo, as static assays fail to capture the dynamic, integrated nature of coagulation in two important ways. First, they overlook the coordinated roles of platelets, fibrin, and VWF in clot formation—interactions that may reveal vulnerabilities only under physiological flow. Notably, botrocetin can induce coagulopathy not just through canonical A1 domain binding, but via secondary mechanisms, including conformational changes in VWF and direct interactions with platelet receptors, disrupting hemostasis even in the absence of traditional binding (Shen et al. 2024). As such, small-bodied opossums with VWF that fails to bind botrocetin in vitro may still exhibit impaired platelet adhesion and fibrin formation under flow—functionally mimicking the response of susceptible species.

Second, shear force—the force per unit area exerted by blood flow along vessel walls—is essential for activating VWF. It initiates the unfolding of VWF multimers, exposing binding sites necessary for platelet capture and clot formation. While large-bodied opossums show resistance to botrocetin at every level—ecological, physiological, and molecular—the same cannot be assumed for small-bodied species. VWF that appears resistant under static conditions may become susceptible under shear, inflating perceived resistance. Thus, in the absence of organismal-level confirmation, it is essential to evaluate VWF–venom interactions under physiologically relevant flow conditions to determine whether small-bodied opossums exhibit true functional resistance.

To address these current gaps in our understanding of venom resistance in small-bodied opossums, we applied a systems-level approach to evaluate multiple physiological responses in *Monodelphis domestica*. Specifically, we tested (1) whether reduced VWF–botrocetin binding translates into functional resistance to coagulopathic effects, and (2) whether this species can inhibit venom components beyond CTLs alone. Using dynamic microfluidic assays, whole blood aggregation, and metalloproteinase inhibition assays, we assessed multifaceted resistance to a range of coagulotoxic venoms (see Graphical Abstract). Because *M. domestica* was also used in prior molecular studies, our design enables direct comparison and extends previous findings into functional, organism-level insight. We find that *M. domestica* is functionally resistant to *Bothrops jararaca* venom across multiple physiological levels, including strong resistance to the integrated coagulopathic effects of botrocetin—the VWF-targeting CTL produced by *B. jararaca*. We also detect species-specific venom resistance across multiple venom protein types, consistent with ecological expectations based on geography and interspecific interactions in this system. Unexpectedly, we discovered that *M. domestica* VWF resists elongation until exposed to much higher shear than human VWF—a striking physiological shift with implications for both venom resistance and human coagulopathies. This work provides the critical physiological evidence that evolutionary signals of venom resistance correspond to robust, organism-level protection in small-bodied opossums, and underscores how studying the molecular and physiological details of natural coevolving systems, especially those involving unexpected or extreme adaptations, can illuminate fundamental insights into basic and translationally relevant physiology.

## Results

### Comparative Microfluidic Tests of VWF elongation, Platelet Adhesion and Fibrin Generation

Under shear, we observed a surprising difference in the dynamics of elongation between opossum and human VWF (Figure 1). Compared to human VWF, opossum VWF required an additional 20 dvn/cm^2^ to fully elongate. In humans, higher molecular weight VWF multimers are more procoagulant than lower molecular weight forms as they are more likely to bind platelets after interacting with subendothelial collagen under flow. Before proceeding, we confirmed there were no obvious differences in multimer distribution between *Monodelphis domestica* and human VWF were observed, with both displaying a typical distribution of high, intermediate, and low molecular weight multimers (Supplementary Figure 1).

**Figure 1.**
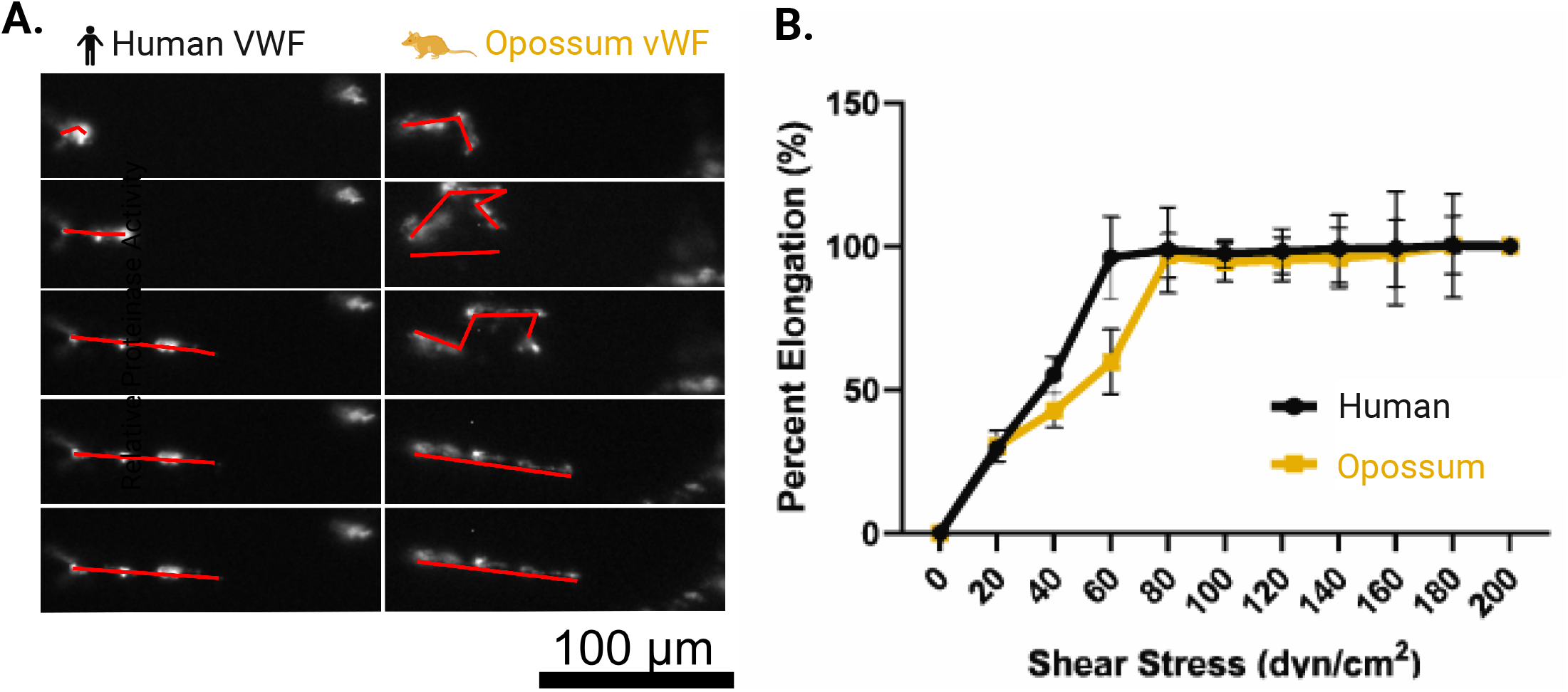
(A) Elongation of von Willebrand Factor (VWF) under different shear conditions. (B)The degree of elongation for both human and *Monodelphis domestica* (opossum) VWF under varying shear forces in a microfluidics assay. Purified VWF was fluorescently tagged and perfused across collagen-coated channels at shear rates of 0 dyn/cm^2^ to 200 dyn/cm^2^ for human and opossum samples, respectively. The percentage elongation of VWF was quantified and normalized to both the 0 dyn/cm^2^ condition (0% elongation) and the 200 dyn/cm^2^ condition (100% elongation). At 60 dyn/cm^2^, human VWF showed full elongation, whereas opossum VWF remained partially coiled, indicating that higher shear rates are required to induce full elongation in the latter. When perfused at 80 dyn/cm^2^, opossum VWF fully elongated, mirroring the response observed in human VWF at 60 dyn/cm^2^. Error bars represent the average ± SD of n = 10 replicates. High degrees of fluorescent labeling caused aggregation or “clumping” of VWF multimers in both human and possum samples, inhibiting complete elongation. Reduced labeling allowed full elongation for human VWF at 60 dyn/cm^2^, but opossum VWF still required the higher shear rate (80 dyn/cm^2^) to achieve comparable uncoiling.

As expected, control experiments with human PRP showed that platelet accumulation on the microfluidic surface was strongly inhibited by botrocetin and convulxin, and weakly inhibited by aspercetin (Figure 2A-D).Venom derived CTLs also significantly reduced platelet fluorescence in *Monodelphis domestica* PRP (Figure 2 E-H); however, the defect was less pronounced after treatment with botrocetin or convulxin than in human PRP (Figure 2 B, F, D, H), suggesting some measure of resistance to those CTLs. In human PRP, the initial rate of platelet adhesion was similar for the control, botrocetin and aspercetin conditions, but reduced to almost zero in the presence of convulxin (as indicated by the initial slope of the platelet fluorescence curves, Figure 2I). A similar trend was observed in opossum PRP, except that platelets adhered even in the presence of convulxin, though at much lower levels than the other conditions (Figure 2J). Concordant with the whole blood aggregometry analyses, the effect of botrocetin on platelet adherence to the microchannel was decreased in *M. domestica* (Figure 2 B, F, J): botrocetin caused a more severe defect in the total extent of platelet adhesion than aspercetin in human PRP, but a less severe defect than aspercetin in opossum PRP (Figure 2I-K). Aspercetin and convulxin caused a similar response in both species (Figure 2K). Concordant with the whole blood aggregometry analyses, the effect of botrocetin on platelet adherence to the microchannel was significantly decreased in *M. domestica* when compared to human (Figure 2 B & F, K). In human PRP, botrocetin caused a 66 ± 16% reduction in the overall extent of platelet adhesion compared to the negative control, whereas it only caused a 39 ± 15% reduction in opossum PRP (Figure 2K).

**Figure 2.**
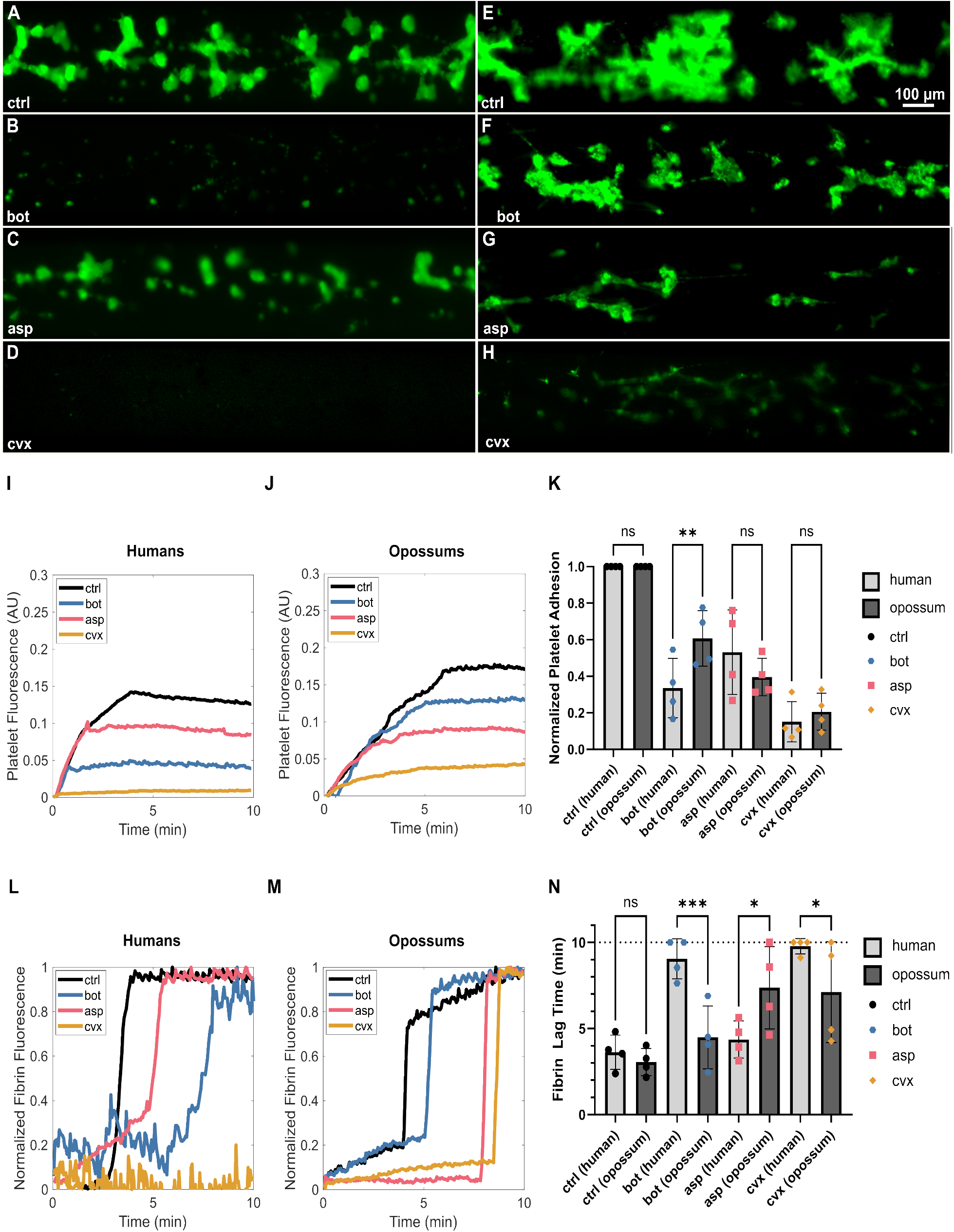
Compared to humans, opossums exhibited resistance to botrocetin (bot), but not aspercetin (asp) or convulxin (cvx). A-H) Endpoint images of platelet (fluorescently labeled in green) adhesion to a collagen coated substrate after 10 min of shear flow at 60 dyn/cm^2^. A) Human platelet rich plasma (h-PFP) perfused across a collagen-coated substrate resulted in robust platelet adhesion in the absence of venom derivatives (ctrl). B) Platelet adhesion was significantly reduced in the presence of bot, C) moderately reduced in the presence of asp, D) and almost completely abrogated in the presence of cvx. Compared to h-PRP, opossum PRP (o-PRP) exhibited a similar extent of platelet adhesion to collagen. However, platelet adhesion was affected to a lesser extent by bot. H) o-PRP responded similar to h-PRP to asp, but H) platelet adhesion in o-PRP was not entirely abolished by cvx. I-J) Representative quantification of platelet fluorescence over time. I) In h-PRP, the overall extent of platelet adhesion was reduced slightly by asp, significantly by bot, and almost completely by cvx. J) Though platelet adhesion in o-PRP was affected by asp to a similar degree as h-PRP, platelet adhesion was only barely reduced by bot. Some degree of platelet adhesion was preserved in the presence of cvx. K) Aggregate platelet adhesion data, normalized to the ctrl condition for each individual donor or opossum. The defect in platelet adhesion was similar for h-PRP and o-PRP for asp and cvx, but reduced significantly in o-PRP for bot. L-M) Representative quantification of fibrin(ogen) fluorescence over time. The sharp ramp indicates fibrin generation, and the corresponding time point is parameterized as fibrin lag time. L) In h-PRP, fibrin polymerized quickly in the absence of venom derivatives, owing to the procoagulant surface of collagen-bound platelets. Asp and bot both delayed the formation of fibrin, with bot having a more pronounced effect. Fibrin did not polymerize during the 10 min experiment in the presence of cvx. M) In o-PRP, bot caused only a marginal prolongation in fibrin lag time. Both asp and cvx significantly delayed fibrin generation. N) Aggregate fibrin generation data. The dashed line at t = 10 min indicates the endpoint of the experiment. On average, fibrin generated in less than 4 min in h-PRP. Bot more than doubled fibrin lag time, with half of the replicates not generating fibrin within the 10 min experimental window. Despite its similarity to bot, asp did not significantly affect fibrin lag time. In three of four replicates, fibrin did not generate within 10 min in the presence of cvx. In o-PRP, fibrin lag time was prolonged by asp and cvx, but not bot. o-PRP generated fibrin faster than h-PRP in the presence of bot or cvx, but slower than h-PRP in the presence of asp. Statistics were performed using one-way ANOVA using Fisher’s LSD correction. Error bars mean ± SD. For significance, *: p ≤ 0.05, **: p ≤ 0.01, *** p ≤ 0.001, ****p ≤ 0.0001.

We further quantified the fibrin generation kinetics (by quantifying ‘lag time’ reflecting the time needed for fibrin clot formation), where a prolonged lag time is unfavorable for effective hemostasis and predictive of bleeding. Purified venom CTLs increased lag time in both humans and opossums (Figure 2 L-N). While convulxin had a dramatic effect in human samples, abrogating fibrin formation almost completely, it showed a more modest response in opossums (Figure 2 L-N). As with platelet adhesion, the greatest signal for resistance was seen in botrocetin treatments (Figure 2 L-N), with an overall reduced effect of botrocetin in opossum PRP (Figure 2 L-N). Botrocetin caused a nearly threefold increase in fibrin lag time from 3.6 ± 1.0 min to 9.0 ± 1.2 min in human PRP, but did not significantly prolong fibrin lag time in opossum PRP (from 3.1 ± 0.8 to 4.5 ± 1.8 min, Figure 2N).

Surprisingly, aspercetin did not significantly increase in the fibrin lag time for human PRP but doubled lag time from 3.1 ± 0.8 min to 7.4 ± 2.4 min in opossum PRP, which is perhaps suggestive of opossum susceptibility to aspercetin. This result is consistent with previously established weak affinity of aspercetin to VWF A1 in protein binding assays and, together with weak platelet adhesion inhibition observed here, suggests that VWF-mediated platelet aggregation is not the primary function of coagulopathy for aspercetin (Drabeck et al. 2022).

Convulxin induced a reduced coagulopathy in opossum PRP compared to human PRP but remained the most potent effector of integrated hemostatic function in both species. Platelet fluorescence and adhesion in response to aspercetin was roughly similar between human and opossum, though aspercetin had a slightly stronger effect on fibrin lag time in opossums. Compared to humans, *M. domestica* VWF showed a measurably reduced integrated coagulopathic response (platelet fluorescence, platelet adhesion, and fibrin lag time) to botrocetin, suggesting that these opossums have functionally meaningful hemostatic protective effects against this CTL.

### Whole Blood Aggregation

We used impedance aggregometry to further probe the species-specificity of opossum resistance to disrupted platelet function. These tests revealed differential, species-specific platelet responses by *M. domestica* to the whole snake venoms and purified CTLs tested (Figure 3). As shown in Figure 3A, collagen, used as a positive control, induced a rapid and sustained increase in impedance, indicative of expected collagen-induced platelet aggregation, while PBS (negative control) established a baseline throughout the 10-minute measurement period.

**Figure 3.**
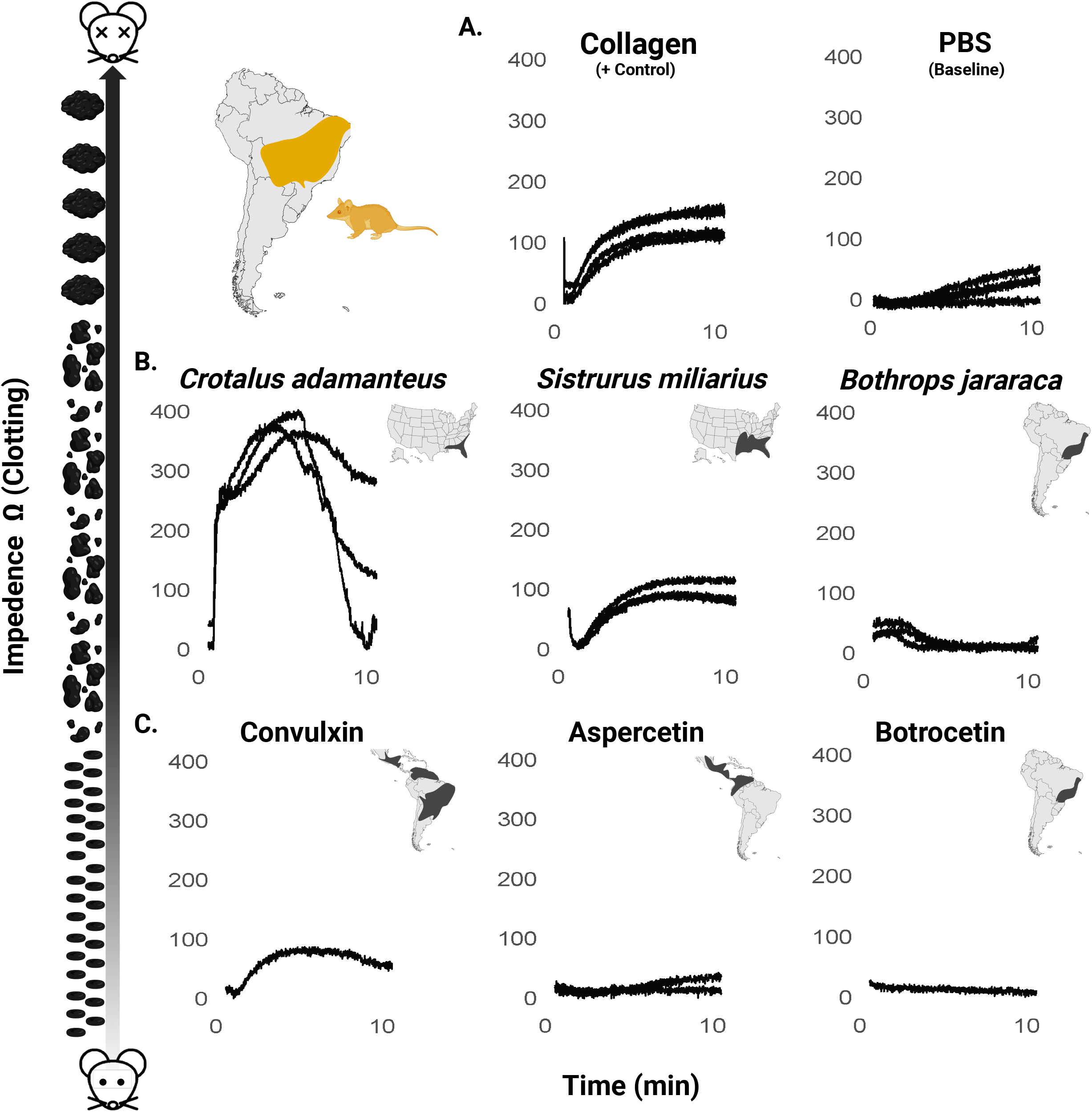
Platelet aggregation *in Monodelphis domestica* whole blood in response to venoms and purified C-type lectins. Impedance was measured over a 10-minute period to assess platelet aggregation response. (A) Collagen (positive control), and PBS (negative control). (B) Whole venoms (*Crotalus adamanteus, Sistrurus miliarius*, and *Bothrops jararaca*) are in the middle panels. (C) C-type lectins, Convulxin (derived from *Crotalus durissus* venom), aspercetin (derived from *Botrhops asper* venom) and botrocetin (derived from *Bothrops jararaca* venom) are on the bottom panels. Data represents the average impedance change over time, normalized to account for baseline variation. Multiple curves represent independent samples (replicates) performed when available. Cartoon on the left depicts platelet aggregation (lowest at y =0 and increases at higher values of y). Range maps of species from which each venom/CTL are derived are shown in the upper right of each graph to allow for a comparison of aggregation response with sympatry.

Among whole venoms, *Crotalus adamanteus* venom elicited the most pronounced platelet aggregation response, with impedance peaking around 5 minutes at approximately 220% of the collagen control (400 Ω) before showing pronounced disaggregation, indicating a robust but transient aggregation effect (Figure 3). *Sistrurus miliarius* venom induced a response reaching approximately 70% of the collagen control (180 Ω), while *Bothrops jararaca* venom resulted in a minimal increase in impedance, remaining just slightly below the PBS baseline (50 Ω), suggesting substantial resistance to platelet activation (Figure 3). As expected, collagen (positive control) produced a peak impedance of 180 Ω, while PBS (baseline control) showed negligible effects with impedance remaining below 50 Ω, which is consistent with standard values for other mammals (Baumgarten et al. 2010; Nash et al. 2017).

The purified CTLs demonstrated variable effects on platelet aggregation. Convulxin, a known collagen-like agonist, induced a notable increase in impedance, mirroring the aggregation pattern of the collagen control but with a delayed onset (Figure 3). Consistent with whole venom results which showed no resistance to *Crotalus* venom, this indicates a lack of resistance to Convulxin (derived from *Crotalus durissus* venom). Aspercetin, which targets VWF, triggered a measurable but extremely reduced platelet aggregation response. Botrocetin, like whole *B. jararaca* venom, elicited no detectable platelet aggregation response. The responses of both aspercetin and botrocetin observed here are extremely minimal compared to previously well-established responses for susceptible species, suggesting strong resistance (Rucavado et al. 2001: Figure 4; Drabeck et al. 2020: Figure 3). Overall, *M. domestica* showed the lowest levels of whole blood aggregation by both snake venom and purified CTLs from their sympatric snake predators in the genus *Bothrops* and appeared to be entirely resistant to both platelet disturbance by *B. jararaca* venom and botrocetin. This is consistent with the hypothesis of evolution of increased resistance to venom of sympatric snakes in *M. domestica*.

**Figure 4.**
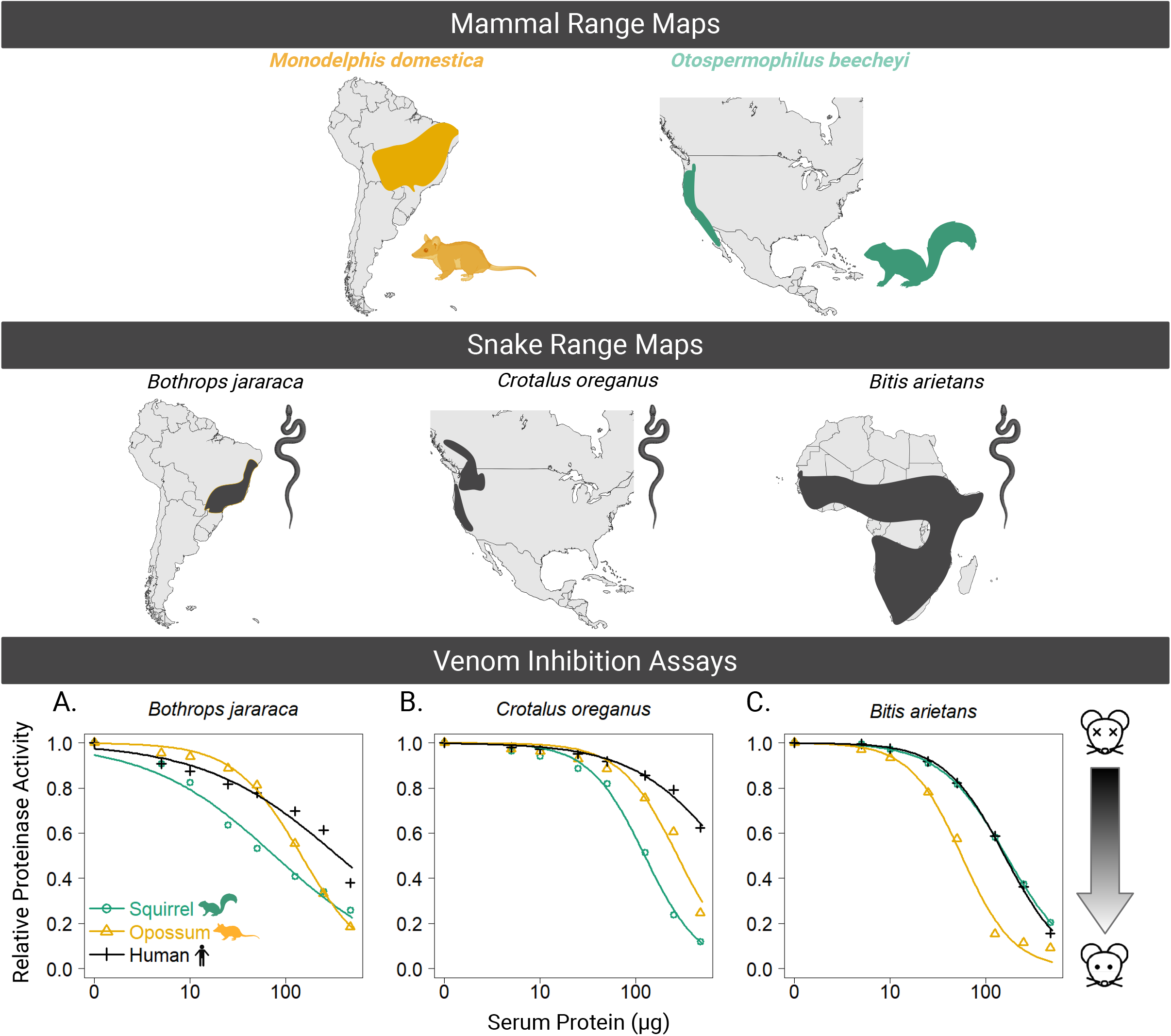
Range maps for mammals *M. domestica* and *O. Beechyi* (top), and snakes (B. jararaca, *C. oreganus*, and *B. arietans* (middle). Bottom panel shows results of the venom inhibition assays. *Bothrops jararaca, Crotalus oreganus*, and *Bitis arietans* venoms were incubated with increasing concentrations of mammalian serum (*M. domestica, O. Beechyi*, and *H. sapiens*), and relative degradation of a gelatin substrate was measured. Data points represent the mean ± SEM for three independent experiments. *M. domestica* serum demonstrated greater inhibition of proteinase activity compared to squirrel and human sera, at higher serum concentrations, particularly for venoms from *B. jararaca* and *B. arietans*. Inhibition was assessed based on the reduction in fluorescence due to the degradation of a gelatin substrate, reflecting the venom’s proteolytic activity. In this graph, the x axis would represent total resistance (depicted with the alive mouse icon), and decrease as you move away from y = 0 (depicted by the dead mouse icon).

### Venom Metalloproteinase Inhibition

Venom metalloproteinases often make up the majority of viper venoms and are thought to be the initial arbiters of venom toxicity, as they degrade vessel walls to release CTLs and other components into the bloodstream(Biardi 2008). Several mammals have coevolved heightened capacity to inhibit metalloproteinases via serum proteins from several families (reviewed in Holding et al. 2016b). We used gelatinase inhibition assays to test the ability of blood serum from different species to inhibit the ability of venom to break down a substrate that mimics the extracellular matrix and are typically used as a proxy quantitation of venom metalloproteinase (SVMP) inhibition. We calculated ED_90_ values, which represent the dosage of serum needed to inhibit 90% of the venom’s gelatinase activity. Blood serum from *M. domestica* demonstrated varying levels of inhibition across the three snake venoms tested: *Bothrops jararaca, Crotalus oreganus*, and *Bitis arietans* (Figure 4A-C; Table 1). In all cases, increasing concentrations of serum protein led to a dose-dependent decrease in venom proteinase activity. For *Bothrops jararaca* venom (Figure 4A), *M. domestica* serum rapidly inhibited SVMP activity at the middle to high concentrations of serum protein tested, resulting in the lowest ED_90_ value of the three sera tested, indicating it inhibited venom approximately 2.5 times better than squirrel serum and 14 times better than human serum (Table 1). Similarly, serum of the squirrel *O. beecheyi* yielded a lower ED_90_ for its sympatric predator, *C. oreganus*, than did the human or opossum sera (Table 1). Finally, opossum serum performed best against the venom of the African viper *B. arietans*, while human and squirrel serum showed overlapping trends in inhibition. Enhanced inhibition for local/sympatric venoms combined with a more general pattern for the African viper venom supports coevolved specificity of mammal serum inhibitors and venom metalloproteinases.

**Table 1.**
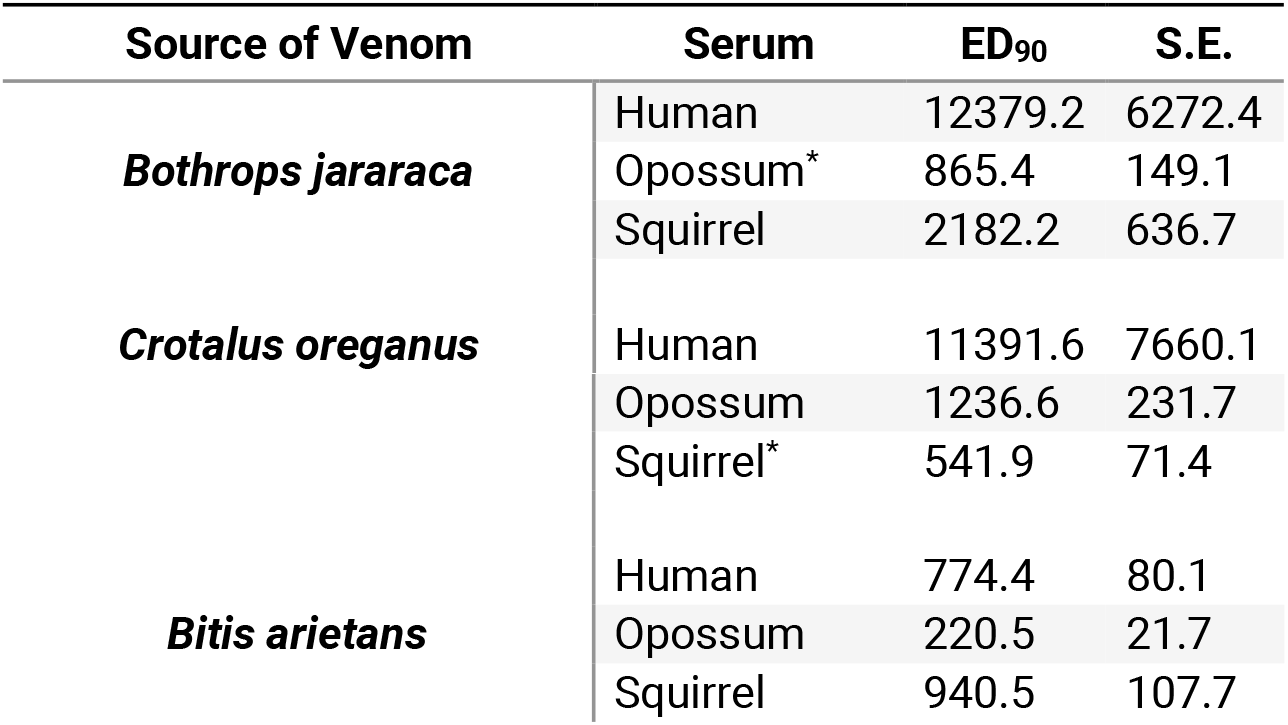
Species-specific ED_90_ values (µg/mL) derived from dose-response analysis of SVMP inhibition by increasing amounts of serum protein from three mammal species. ED_90_ (Effective Dose, 90%) indicates the amount of serum required to inhibit 90% of the venom’s proteinase activity. Asterisks indicate species that are sympatric with the snake venom tested.

## Discussion

Understanding the physiological mechanisms, molecular underpinnings, and evolutionary history of venom resistance across opossums is crucial for understanding how selective pressures, ecological interactions, and physiological constraints shape the evolution of such extreme adaptations (Byrne et al. 2014; Xu et al. 2020). Our results provide compelling evidence that *M. domestica* possesses considerable resistance to both CTLs and whole venom from *Bothrops jararaca*, and suggests that previous *in vitro* work showing convergently derived mutations on VWF is indicative of realized venom resistance (Drabeck et al. 2022). We also uncover a previously undescribed modification to opossum VWF, that it requires a greater shear force to elongate, potentially contributing to the resistance phenotype or other important adaptive physiologies.

Under normal conditions, VWF will be induced to elongate and bind platelets in response to either exposed collagen (mimicking vessel damage) or under increased shear force (caused by increased blood flow after vessel damage), creating a primary platelet plug preceding the formation of a more stable cross-linked fibrin clot (Sadler 1998; Ruggeri 2003). Botrocetin is known to induce coagulopathy upon introduction to the body by binding to VWF (at the A1 domain) and inducing VWF to elongate and attach to platelets (Matsushita et al. 2000; Fukuda et al. 2002). Botrocetin then changes conformation slightly and binds secondarily to the platelet GP1Bα binding site, creating a tight trimolecular complex which renders platelets unavailable for adhesion and causes extreme thrombocytopenia (Fukuda et al. 2002, 2005). More recently, detailed physiological work on botrocetin has shown that in the presence of VWF, botrocetin can induce coagulopathy even in the absence of its preferred platelet receptor (GP1Bα), and that it also interacts directly with a different platelet integrin (αIIbβ3) to block fibrinogen and other ligands essential for coagulation and healing (Shen et al. 2024). While aspercetin is also a VWF-mediated agonist from a closely related viper (*Bothrops asper*) and is thought to work in a similar manner because of its ability to induce VWF-mediated platelet agglutination, detailed structural studies have yet to locate its exact binding site on VWF and prior kinetics studies show weak binding to the A1 domain (Rucavado et al. 2001; Drabeck et al. 2022). Integrated coagulopathic data shown in this work supports that VWF-mediated platelet activity is a weak action for aspercetin and may not be the main mode of coagulopathy for this CTL. Convulxin is derived from the more distantly related tropical rattlesnake *Crotalus durissus*, and binds directly to the GPVI platelet receptor, inducing coagulopathy independent of VWF (Batuwangala et al. 2004).

Our assays of whole blood aggregation revealed a reduced response of *M. domestica* blood to both botrocetin and aspercetin but not convulxin. Initially, this might lead us to assume that opossum resistance is specific to CTLs from *Bothrops spp*. and centered at VWF. However, microfluidic assays showed that platelet adhesion and fibrin clot formation in opossums was more disrupted by aspercetin than botrocetin. This suggests that the molecular changes that have previously been found to abolish binding between opossum VWF and botrocetin result in resistance to this CTL (implying specificity), but *does not* result in resistance to aspercetin, despite the appearance of strong resistance that would be gleaned from aggregation data alone. Aspercetin appears to be able to function nearly as well in opossums as it does in humans in microfluidics assays (in both cases it has weak function), but not in whole blood assays, highlighting the importance of integrated assays which capture a more complete picture of coagulopathic function. These results suggest that aspercetin may act directly on platelets, fibrin, or other agonists more strongly during platelet adhesion and fibrin clot formation, or that its function in opossum depends on the presence of shear force. Given that its action in humans is also weak relative to other CTLs, and previous work has revealed a surprisingly low affinity for human VWF compared to other VWF-mediated agonists, it is possible that this is an accessory function of this protein, or that its affinity for VWF is specific to a non-mammalian VWF target.

Similarly, we observed a tremendously decreased response in opossums (compared to previously established human responses) to whole venoms from both species of *Bothrops*, but not *Crotalus*. Opossum serum showed superior ability to inhibit the enzymatic action of whole venom for all species tested, except *Crotalus oreganus*, which was best inhibited by its sympatric prey (*Otospermophilus beecheyi*). Interestingly, *M. domestica* serum very strongly inhibited venom from the African viper, *Bitis arietans*, which points to the likelihood that this species also possesses generalized inhibitory proteins (snake venom metalloproteinase inhibitors, phospholipase inhibitors), similar to those seen in other highly resistant taxa (Voss and Jansa 2012).

*Bothrops spp*. are well known to interact with opossums and are a known prey item of several species of South American opossums (Voss and Jansa 2012; Voss 2013). *Bothrops jararaca* co-occurs across the range of *M. domestica*, are relatively small and gracile, semi-arboreal, and have been anecdotally observed to bite *M. domestica* without effect (Martins et al. 2001; Da Silva et al. 2024; exclusivo cuíca predando cascavel e coral verdadeira e é picado por elas n.d.). *Bothrops asper* are somewhat closely related to *B. jararaca;* however they are much larger and their range does not overlap with *M. domestica*. Previous work examining diet in small-bodied opossums has suggested that predator-prey relationships between opossums and snakes may role-switch relative to factors affecting venom potency like geographic region, body size, and age (Voss 2013). Conversely, *Crotalus durissus* is documented to be prey of only the larger bodied opossums (Didelphini) and though its range overlaps with *M. domestica*, our data did not reveal compelling evidence that *M. domestica* has profound resistance to convulxin (the CTL derived from its venom) (Almeida-Santos et al. 2000; Voss and Jansa 2012). This suggests that small-bodied opossum venom resistance may be more specialized for species that they target for prey, are appropriately sized for prey, or that are more abundant in their environment.

Interestingly, our findings reveal that VWF in *M. domestica* requires a comparatively higher shear force than that for human VWF, to elongate. This property could have important implications for venom resistance by making it more difficult for venom components, like botrocetin, to bind VWF and exact subsequent coagulopathic effects. Elevated shear thresholds for VWF elongation may also protect against other physiological disruptions, such as spontaneous platelet adhesion in high-flow conditions or inappropriate clot formation. Beyond venom resistance, this trait may be linked to broader physiologies of non-placental mammals, potentially reflecting differences in vascular dynamics, metabolism, or reproductive strategies. The discovery of this and potentially other species with shear-resistant VWF may also provide a novel avenue for therapeutic exploration, where engineering similar shear-resistant VWF variants could help prevent bleeding disorders associated with left ventricular assist devices (LVAD) (Nascimbene et al. 2016). While there is some limited data to suggest that increased shear elongation may occur in other venom resistant species (pigs), little is known about the diversity of this trait across species, or what potential costs and tradeoffs it might evoke (Chan et al. 2017). Further work is merited to better understand this unique physiology and explore its potential for inspiring new strategies for mitigating LVAD-related complications and other vascular disorders.

Our work adds detailed comparative assays using physiologic vessel models to demonstrate that venom resistance is present in small-bodied opossums, and highlights the ability of comparative analyses to inform otherwise cryptic adaptations and ecologies (Jansa and Voss 2011; Drabeck et al. 2020, 2022). It is likely that many more of the opossum species identified to have similar and convergently evolved changes in VWF may have similar degrees of venom resistance (Drabeck et al. 2022). In light of this, further work is warranted to examine the true ecological roles and behaviors of this entire group of mammals. These results also provide vital insight into the evolution of venom in this region and suggests that South American viper venoms may have evolved in the presence of many more species of co-evolving venom-resistant mammalian predators and prey than previously assumed. Detailed physiological data presented here serves to further inform the physiological mechanisms, relative strength, and specificity of this extreme adaptation, and provides key insights for future work aiming to understand the molecular basis and evolution of this remarkable trait.

## Methods

### Venom and Protein Purification

Purified fractions of botrocetin from *Bothrops jararaca* and aspercetin from *Bothrops asper*, bitiscetin from *Bitis arietans*, as well as whole venom from *Bothrops jararaca* were used from previous work (Drabeck et al. 2020, 2022)(botrocetin B). Convulxin was purchased from ChemCruz (Santa Cruz Biotechnology Inc, Santa Cruz, CA). Whole venom from *Crotalus oreganus* and *Sistrurus miliarius* was obtained during previous studies (Gibbs et al. 2011). Venom CTLs were chosen to best provide a direct comparison of physiological and protein assays done in previous work (Drabeck et al. 2020, 2022).

### Opossum Handling and Blood Collection

Opossums (n= 16) were housed in the Oberlin research colony and cared for according to procedures previously described (Rousmaniere et al. 2010). To collect blood for VWF purification, opossums (*Monodelphis domestica*) were sacrificed by first being placed under isoflurane sedation and then exsanguinated via the posterior vena cava, per IACUC protocol F23BBYC-1 (Keyte and Smith 2008). Samples used for VWF purification were collected in an ACD-coated syringe (acid citrate dextrose Solution A) and preserved in vacutainers containing the same solution (BD Pharmaceuticals). Samples used in all other assays were collected in 3.2% sodium citrate coated syringe and similarly placed in vacutainers containing the same solution (BD pharmaceuticals). Human blood was collected from de-identified donors via venipuncture into 3.2% sodium citrate vacutainers (BD Pharmaceuticals) by the Case Western Reserve University Hematopoietic Biorepository and Cellular Therapy Core under an Institutional Review Board (IRB)-approved protocol (Case 12Z05, IRB # 09-90-195) of University Hospitals Cleveland Medical Center. Blood was then centrifuged at 150 rcf for 15 min at 25°C to isolate a platelet-rich plasma (PRP) supernatant, which was removed without disturbing the buffy coat. The remaining sample was then centrifuged at 2000 rcf for 25 minutes to recover platelet-poor plasma which was later used for dilutions and standards.

### Microfluidic Assays

We performed microfluidics assays developed to approximate *in vivo* injury conditions which allowed for real-time measurement of VWF/VWFplatelet recruitment and fibrin generation. Microfluidics experiments were performed in the Bioflux 200 system (Cell Microsystems) using 48-well High Shear Plates (Cell Microsystems) (Conant et al. 2009; Girish et al. 2022). Glass-bottomed microfluidic channels with a width of 250 µm and a height of 70 µm were coated with 100 µg/mL type 1 equine fibrillar collagen (Chrono-log Corporation) in 20 µM acetic acid for 60 min at 37°C. These conditions facilitate the self-polymerization of collagen fibers which anchor to the glass substrate, creating an environment which mimics collagen exposure following vascular injury. Unbound collagen was washed by perfusing 0.1 wt% bovine serum albumin (BSA) in calcium-free phosphate buffered saline (PBS) at 30 dyn/cm^2^ for 2 min.

While we have previously standardized this assay for use with human blood, it was unknown whether the equine collagen would allow binding of opossum VWF in a similar manner and under similar shear flow conditions. To verify that this assay is applicable to opossum VWF, we used purified and fluorescently tagged VWF to determine experimental conditions and verify that opossum VWF binds effectively to equine fibrillar collagen used in standard microfluidics assays (Figure 1). AF488-labeled human or opossum VWF in calcium-free PBS was perfused through the collagen-coated microfluidic channel at shear stresses ranging from 20 to 200 dyn/cm^2^. VWF binding to collagen was imaged under epifluorescence using a Zeiss Axio Observer microscope with a 10x objective (Plan-NEOFLUAR, NA = 0.3). We observed similar overall VWF deposition, indicating that opossum VWF binds to equine collagen to a similar extent as human VWF. We further observed that opossum VWF required comparatively higher shear flow conditions (80 dyn/cm^2^ for opossum VWF vs. 60 dyn/cm^2^ for human VWF) to fully elongate, which we discuss in detail in the results and elsewhere (Figure 1).

With assay conditions established, we assessed the ability of *M. domestica* to resist the detrimental effects of venom CTLs. Opossum or human (as a venom non-resistant control species) blood was processed and used to perform assays within four hours of any given draw, to ensure that platelet activity in the blood remained viable. Citrated PRP was obtained as described in the *Opossum Handling and Blood Collection* section. PRP was incubated with 1 µg/mL calcein-AM (Invitrogen) to stain platelets and 10 µg/mL AF-647 fibrinogen (Invitrogen) for 30 min at room temperature. Samples were recalcified to 20 mM CaCl_2_ and immediately perfused through collagen-coated channels at 60 dyn/cm^2^, a shear condition that is relevant to wound environments. These conditions provide more fine-tuned information about the degree and type of physiological response induced by exposed collagen in the presence of agonists. Epifluorescence microscopy was used to image platelet accumulation and fibrin generation in real time. Samples were run without any additives (Control) to assess normal platelet adhesion and fibrin generation. For each sample, complementary assays were performed in the presence of three venom derived CTLs known to affect platelet aggregation function: botrocetin, aspercetin, and convulxin. Each CTL was used at its ED_90_ concentration for human blood: 4 µg/mL for botrocetin, 20 µg/mL for aspercetin, and 0.5 µg/mL for convulxin. In this assay, we expect that CTL-induced VWF/platelet complexation leading to thrombocytopenia and low VWF levels should result in reduced platelet adhesion and prolonged fibrin generation times, mimicking bleeding observed *in vivo*. To account for differences in platelet counts between individual animals, platelet adhesion was normalized to the control condition for each sample.

### Aggregation Assays

To test *M. domestica*’s ability to resist whole venom as well as isolated CTLs, whole blood aggregation was assessed using an impedance aggregometer (Roche multiplate aggregometer). Briefly, these assays differ from microfluidic assays as coagulopathic response is indicated by *induction* of platelet aggregation, rather than inhibition. In the absence of exposed collagen and/or shear force, platelets in blood (in a stationary cuvette) should not rapidly aggregate unless exposed to a platelet activating agonist (i.e. whole venom or pure venom-derived CTLs). Failure of agonists to induce platelet aggregation suggests strong venom resistance to coagulopathic venom components, including CTLs. Thus, we expect *M. domestica*’s platelet aggregation response to be lower for sympatric venom and CTLs than for allopatric venom and CTLs, suggesting higher resistance to species they have evolved with.

Platelet aggregation response in this assay is measured by impedance, where higher impedance means a greater response. Purified CTLs used were aspercetin (20 µL, 0.1 mg/mL), convulxin (20 µg, 0.01 mg/mL), and botrocetin (20 µg, 0.01 mg/mL). Platelet aggregation was measured over 10 minutes, with impedance changes recorded as a measure of platelet aggregation. All purified CTLs were selected from South American vipers that were either completely sympatric (*Bothrops jararaca*), partially sympatric (*Crotalus durissus*), or allopatric (*Bothrops asper*) with *M. domestica*. It should be noted that while aspercetin and botrocetin induce aggregation via VWF, convulxin induces aggregation via a direct interaction with the platelet. In addition to purified CTLs, we also challenged *M. domestica* with whole venom from the same sympatric species (*Bothrops jararaca*), as well as two distantly related allopatric species with vastly diverse venoms (*Crotalus adamanteus* and *Sistrurus miliarius*) to provide some measure of the specificity/cross-reactivity of venom resistance in *M. domestica* in the absence of availability of whole venom from *Bothrops asper* and *Crotalus durissus* (see Figure 3 range maps).

Venous blood samples collected from *M. domestica* in 3.2% sodium citrate tubes were tested within four hours of collection at room temperature. For each assay, 300 µl citrated whole blood and 300 µl NaCl/CaCl2 solution (0.9% saline and 3mM CaCl2 provided by the manufacturer) were aliquoted into cuvettes and stirred at 37°C. Aggregation was induced by adding 20 µL of agonists to the samples. Collagen (20 µg/mL) was used as a positive control, PBS (phosphate buffered saline) was used as a negative control, Aggregation response was calculated as the average of two replicate measures. To facilitate cross-comparison, the data were normalized by subtracting the minimum value within each treatment group, ensuring that the minimum impedance value was zero. Data visualization was performed using R software to allow for comparison of platelet responses to different agonists. Venom assay and controls were conducted in triplicate to ensure reproducibility; however, due to limited plasma volume, CTL tests were not replicated.

### VWF Multimer Gel

Opossum VWF was purified by size-exclusion chromatography at 4°C as previously described (De Marco and Shapiro 1981), using plasma pooled after high-speed centrifugation. Fractions were quantitated by absorbance at 280 nm using the in-line detector on the AKTA Pure. To ascertain that VWF from *M. domestica* bound to microchannel equine-derived type 1 fibrillar collagen (Chrono-log Corporation), purified and quantified opossum VWF was tagged with Alexa488 using the Alexa Fluor 488 (AF488) Microscale Protein Labelling Kit (Invitrogen) with minor modification for subsequent microchannel runs. The remaining untagged VWF was frozen and stored at -80C.

To compare opossums’ VWF multimer structure to human, we performed a multimer analysis using VWF purified from *M. domestica* plasma and HumateP (a purified human plasma-derived product that contains VWF and Factor VIII, CSL Behring). Due to the enormous size of intact VWF multimers, agarose is used instead of polyacrylamide. VWF multimers were analyzed by electrophoresis using a 1.6% separating LDS agarose gel and visualized using the rabbit anti-human VWF antibody A0082 (DAKO) as previously described (Pruss et al. 2011). Protein was transferred to nitrocellulose using the semi-dry Power Blotter (ThermoFisher) and western blot was performed using the iBind Flex (Invitrogen) system with the rabbit anti-human VWF antibody A0082 (DAKO) as primary and IRDye 800CW anti-rabbit antibody (Li-Cor). Lanes were visualized a Li-Cor Odyssey CLx Imager and Image Studio (LiCor).

### Venom Inhibition Assays

We evaluated the inhibition of whole venom by *Monodelphis domestica* serum against three snake venoms (*Bothrops jararaca, Crotalus oreganus*, and *Bitis arietans*) using a modified protocol from the EnzChek Gelatinase/Collagenase Assay Kit (Life Technologies, Carlsbad, CA, USA). Briefly, this assay tests the ability of *Monodelphis domestica* serum to neutralize venom-induced degradation of a gelatin substrate, which mimics the extracellular matrix proteins targeted by snake venom metalloproteinases (SVMPs), and other enzymatic venom components (Holding et al. 2016a). The DQ-gelatin substrate was diluted to 1:20 in reaction buffer for the assay.

Serum was taken from opossums during venous blood extraction and frozen for subsequent analyses, and the protein concentration of each serum sample determined with the Pierce BCA Protein Assay Kit. Venom samples were diluted to a final protein concentration of 0.3125 ng per well. Venom was incubated with various amounts of *M. domestica* serum, with 0, 5, 10, 25, 50, 125, 250, or 475 μg serum protein per well, to determine the extent of venom inhibition.

Fluorescence intensity was measured in relative fluorescence units (RFU) using a SpectraMax M2 microplate reader (Molecular Devices) at 30 second intervals. The slope of the fluorescence increase (RFU min^− 1^) from the linear portion of the reaction curve was calculated to quantify the venom’s SVMP (snake venom metalloproteinase) activity. Decreased slope values, as compared to control reactions with no serum, indicate successful inhibition of venom enzymatic activity by the serum. Each experiment was performed in triplicate, and data were normalized to the control (untreated venom) to calculate relative SVMP activity.

Dose-dependence curves were fitted using the drc package in R, with a four-parameter logistic model for each serum treatment (opossum, squirrel, and human) (Ritz et al. 2015). Human plasma was used as a non-coevolved venom-sensitive baseline for venom SVMP inhibition by serum proteins, and squirrel (*Otospermophilus beecheyi*) was used to compare species-specific inhibition as it has previously been shown to have resistance for its sympatric predator (*Crotalus oreganus*) (Holding et al. 2016a). Venoms were selected to represent 1) a sympatric species (*Bothrops jararaca*) for which ecological and other data predict resistance, 2) an allopatric species (*Crotalus oreganus*) for which another mammal (ground squirrels) have evolved venom resistance 3) an allopatric distantly related viper (*Bitis arietans*) (Figure 4 top and middle panels).

## Acknowledgements

The authors would like to thank Alexandra Rucavado, Erika Hingst-Zaher, and the Kentucky Reptile Zoo for their help in acquisition and purification of venoms used for this work. Authors on this work were supported by the Minnesota IRACDA Program 2K12GM119955 (NIGMS, NIH), the NSF STEM-APWD (DEB 2316784), NIH R01-HL121212, NIH R35-HL171421, and NIH R35-HL150784 (NHLBI). The authors would like to thank Suzanne McGaugh for tremendous guidance as well as lab and resource support. The authors would also like to thank Marvin Neiman (Case Western), Alison Narayan/Sarah Ackenhusen (University of Michigan), and Randy Westrick (Oakland University) for their guidance, feedback, and support.

## Data Availability

All raw data generated in this study, including microfluidic assay results, whole-blood aggregation data, and gelatinase inhibition assay outputs, including scripts for analysis and visualization are publicly available via the Zenodo Digital Repository: 10.5281/zenodo.15285263.

## Conflict of Interest Statement

Matthew Holding, Dante Disharoon, Laura Haynes, Bipin Chakravarthy Paruchuri, M. Hao Hao Pontius, Krista Golden, Jordan A. Shavit, Karl Desch, David Ginsburg, Anirban Sen Gupta, Yolanda Cruz, and Danielle H. Drabeck declare that they have no known competing financial interests or personal relationships that could have appeared to influence the work reported in this paper.

**Supplementary Figure 1.**
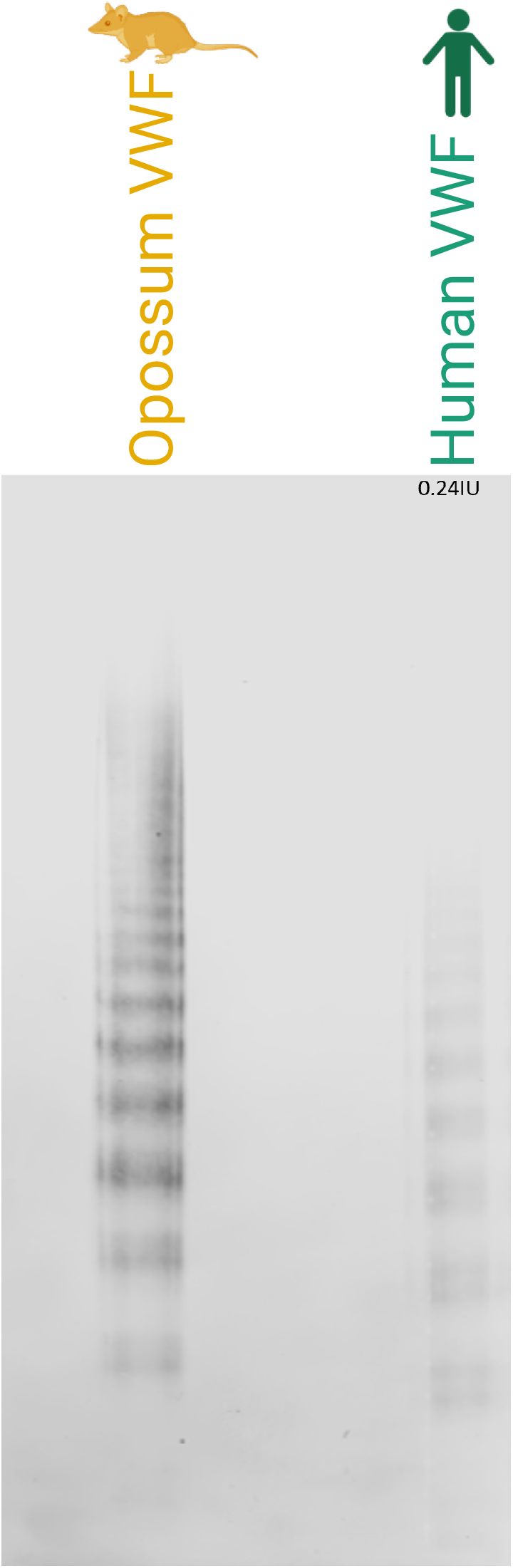
Opossum VWF Multimer Gel 1.6% resolving + 0.8% stacking Tris Glycine LDS-SeaKem HGT Agarose Vertical Multimer Gel (8×8cm vertical). Multimeric structures of von Willebrand factor (VWF) were analyzed using a 1.6% lithium dodecyl sulfate (LDS) agarose gel under non-reducing conditions. Lane 2 contained 30 µL of purified Monodelphis domestica VWF (heated at 60°C for 30 minutes) and 10µl loading buffer. Lane 5 contained 30 µL HumateP (human VWF) as a control (heated at 60°C for 30 minutes) and 10µl loading buffer. The gel was transferred to a nitrocellulose membrane and visualized by fluorescence with anti-human VWF antibody (Dako A0082) using a Li-Cor Odyssey CLx Imaging System (Li-Cor).

