## Supplementary figures and images for "Species-specific, multifaceted venom resistance in *Monodelphis domestica* reveals novel physiological behavior of von Willebrand Factor under flow"

### Supplementary Figure 1

#### Supplementary Figure 1

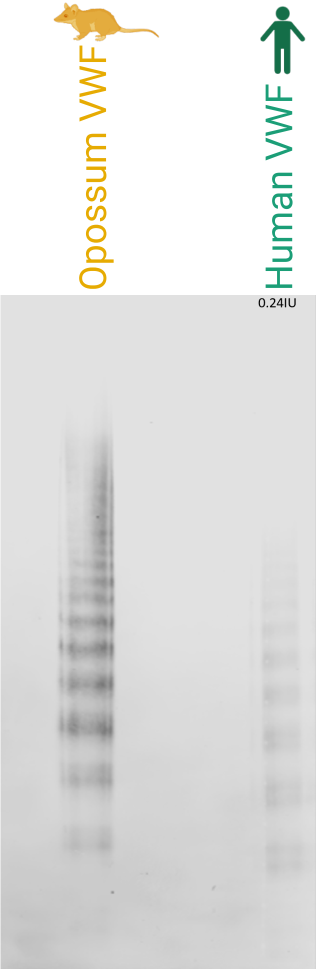
